# Photosynthetic adjustments maintain lettuce growth under dynamically changing lighting in controlled indoor farming setups

**DOI:** 10.1101/2025.01.16.633207

**Authors:** Arttu Mäkinen, Hirofumi Ishihara, Sylvain Poque, Nina Sipari, Kristiina Himanen, Ilona Varjus, Juho Heininen, Matti Pastell, Paula Elomaa, Alexey Shapiguzov, Titta Kotilainen, Saijaliisa Kangasjärvi

## Abstract

Studies have uncovered delicate mechanisms that enable plant acclimation to fluctuating light. Translating the knowledge to controlled environment agriculture could advance the development of cost-effective dynamic lighting strategies, but the effects of varying light intensities on vegetable crops remain poorly understood. Here we recorded chlorophyll fluorescence, photosynthetic activity, metabolic responses, and growth of lettuce (*Lactuca sativa* L.) cv. ‘Katusa’ under dynamic lighting. The light intensity was varied at different times of the photoperiod with uniform daily light integral. Three setups, including a plant phenotyping facility, a small-scale vertical farm testbed and a larger-scale vertical farm were utilized to address the physiological responses and scalability of lighting strategies. We found that dynamic lighting supported lettuce cv. ‘Katusa’ growth in all three indoor cultivation setups, even under artificial “split-night” regimes where the photoperiod was interrupted by two periods of darkness. The lettuce plants displayed delicate adjustments in photosynthetic light reactions and carbon metabolism, the latter of which followed the cumulative daily light integral under different lighting regimes. However, the overall metabolic composition of lettuce leaves did not respond to the changing light intensities. Our findings support the conclusion that dynamic lighting enables cost-effective lighting via optimization of electricity use in indoor cultivation.

**Highlight:** Photosynthetic adjustments maintain lettuce (*Lactuca sativa* L.) cv. ‘Katusa’ growth under dynamic lighting. This enables cost-effective cultivation via optimization of electricity use in controlled environment agriculture.

## Introduction

Light is an important environmental factor that affects the productivity and nutritional value of crops. In natural conditions, wind-induced canopy movements, cloudiness, and climatic and seasonal alterations affect the light environment at different time scales (Kaiser et al., 2019, Slattery et al., 2018, Kotilainen et al. 2020). Plants respond to changes in the direction, duration, intensity, and spectral quality of light, which in natural conditions alternate seasonally and within a 24-hour diurnal cycle (Brelsford et al., 2019, Sellaro et al., 2010), and are affected by neighbouring plants (Ballaré et al., 1987, Casal, 2013). Studies have uncovered fast photosynthetic rearrangements and more durable light-induced metabolic and developmental adjustments, which may affect plant productivity (Morales and Kaiser, 2020, Hotta, 2021, Dantas et al., 2023, Poorter et al., 2019, Slattery et al., 2018, Kaiser et al., 2019). More recently, application of changing light conditions has been recognized as a way to achieve cost-effective lighting in controlled environment agriculture (CEA) (Kaiser et al, 2024, Bechtold et al, 2024, Lazzarin et al, 2024). In dynamic lighting strategies, the intensity of artificial lighting is purposely alternated during indoor cultivation.

Dynamic adjustment of lighting can be achieved with light-emitting diode (LED)-luminaires fitted with adjustable light intensity output capabilities that enable dimming. An intrinsic property of LEDs is that their light intensity output is dependent on the electrical current. Therefore, by regulating the supplied electrical current, their intensity and electricity consumption can be adjusted almost instantaneously. When paired with modern control and automation systems, dimmable LED-luminaires allow adjustment of the light environment in indoor cultivation. Ability to dynamically adjust lighting is becoming increasingly important, since lighting and heating make up a significant part of the production costs in CEA, especially in wintertime. European power markets have changed in recent years with record-breaking levels and fluctuations in prices (Cevik & Ninomiya 2022). The Nord Pool power exchange (https://www.nordpoolgroup.com/) publishes hourly spot electricity prices daily, 24 hours in advance, which would allow optimization of light intensity in CEA according to the electricity price. However, how varying light cycles affect crop quality and productivity remain insufficiently understood.

In photobiological studies, analysis of chlorophyll fluorescence enables non-invasive assessment of photosynthetic performance. In photosynthetic light reactions, electrons flow from Photosystem II (PSII), cytochrome b6f complex (Cytbf) and photosystem I (PSI) to NADP^+^, generating reducing power in the form of NADPH. The electron transfer reactions also generate a proton motive force across the thylakoid membrane, which drives the production of ATP by the ATP synthase. In photosynthetic carbon metabolism, a variety of enzymes consume NADPH and ATP for carbon fixation, starch biosynthesis, and other biosynthetic processes (Stirbet et al., 2020). In a changing light environment, transient perturbations in photosynthetic electron transfer reactions may enhance photoinhibition of PSII, commonly measured as the chlorophyll fluorescence parameter Fv/Fm (Baker, 2008, Murchie and Lawson, 2013, Järvi et al., 2015, Li et al., 2018). At the same time, acidification of thylakoid lumen activates dissipation of excess excitation energy as heat at the light-harvesting complex of PSII (LHCII), which can be measured as increased qE, the major component of non-photochemical quenching of chlorophyll fluorescence (NPQ) (Niyogi et al., 1998, Ruban, 2016, Walter and Kromdijk, 2021). On a cellular level, chloroplast movement can optimize light harvesting, albeit the physiological role of light-induced chloroplast relocation has recently been debated (Wilson and Ruban, 2020).

Photobiological studies on the model plant *Arabidopsis thaliana* have shown that photosynthetic activities are sensitive to the frequency, duration, and magnitude of light intensity changes (Vialet-Chabrand et al., 2017, von Bismarck et al., 2023). Matthews et al (2018) found that photosynthetic gas exchange responded to the intensity and pattern of changing light intensities, and that the responses varied at different times of the day. Notably, reduced accumulation of biomass was a common response to stressful changes in light intensities, when compared to *A. thaliana* rosettes grown under constant light intensity (Morales and Kaiser, 2020).

Among commercially cultivated leafy vegetables, lettuce (*Lactuca sativa* L.) was reported to tolerate moderate light intensity changes without negative effects on growth, whereas more drastic, ten-fold changes in light intensity impaired photosynthetic activity and growth (Bhuiyan and van Iersel, 2021). While the underlying molecular mechanisms remain unresolved, crop performance under different frequency, duration, intensity, and spectrum of dynamic lighting should be examined to avoid stress-induced growth reduction and formation of off-flavour in commercial produce (Kaiser et al, 2024).

Here we analysed the physiological performance and growth of lettuce cv. ‘Katusa’ under dynamic lighting. Three different cultivation setups, including a small-scale vertical farm testbed, a plant phenotyping facility, and a larger-scale experimental vertical farming system were utilized to address the physiological responses and scalability of the dynamic lighting strategies. The light intensity was varied at different times of the photoperiod, with uniform daily light integral (DLI) across different lighting regimes within each cultivation set-up. We find that lettuce responds to dynamic lighting by delicate adjustments in photosynthetic light reactions and carbon metabolism, while the overall metabolite profiles remain largely unaltered. Our findings suggest that application of dynamic lighting offers cost-effective cultivation solutions that support lettuce growth without impairing its nutritional quality in controlled indoor farming setups.

## Materials and methods

### Plant material and growth conditions

Lettuce (*L. sativa* L. cv. ‘Katusa’) seeds were obtained from Puutarhaliike Helle Oy, Lieto, Finland. The plants were grown in three different experimental set-ups, including a small-scale vertical farm testbed constructed in-house, National Plant Phenotyping Infrastructure (NaPPI; https://www.helsinki.fi/en/infrastructures/national-plant-phenotyping, PlantScreen™ Compact System, Photon Systems Instruments, PSI, Drásov, Czech Republic), and a larger-scale vertical farming system (VIS, Vacuum Insulation Solutions Oy, Espoo, Finland), as detailed in Supplemental Table 1.

In experiments performed in a small-scale vertical farm testbed, pots with stratified seeds were distributed to irrigation trays that were divided to four light-isolated cultivation shelf compartments. For nine days of initial growth, the conditions were set as follows: light intensity of 155 μmol m^-2^ s^-1^ photosynthetic photon flux density (PPFD), photoperiod of 18 h light and 6 h dark (L18:D6), air temperature during the day 21°C, and night 19°C, and relative humidity (RH %) of ∼40%. The positions of the pots were randomly rearranged multiple times during the experiments to alleviate possible border effects and environmental gradients.

In the phenotyping experiment, pots with stratified seeds were distributed to 24 imaging trays positioned in eight light-isolated cultivation shelf compartments (CS 250/300_4_2.4, PSI, Drásov, Czech R.). For 12 days of initial growth, the conditions were as follows: light intensity 155 μmol m^-2^ s^-1^ PPFD, photoperiod L18:D6, air temperature during the day 21°C, and night 19°C, and relative humidity 60-70%.

The larger-scale vertical farming system set up was located at the Natural Resources Institute Finland research station in Piikkiö, Finland. The seeds were germinated in darkness in a climate controlled dark room for 1 day at 18°C and thereafter transferred to a seedling line in a standard greenhouse under automatically controlled climatic conditions (Itumic, Jyväskylä, Finland). The temperature was set to 18°C, and the relative humidity fluctuated between 60 and 80%, 800 ppm of CO_2_ was supplied. Light intensity was 350 μmol m^-2^ s^-1^ PPFD, with photoperiod L18:D6. After two weeks of initial growth, 264 plants per experiment (144 for imaging and 120 for final harvest measurements) were transferred into a hydroponic nutrient film technique-cultivation (NFT) system located in the larger-scale vertical system divided into three light-isolated compartments.

### Lighting regimes and spectral composition

Adjustable (0–100%) and independently programmable digital LED-light controllers connected to dimmable LED-lights were used to implement the varying light intensity regimes in all three experimental setups. In the plant phenotyping experiment, LED-light controllers embedded in NaPPI cultivation shelves (LC, PSI, Drásov, Czech R.) were used, whereas in the small-scale vertical farm testbed and the larger-scale system, universal LED-light control system (GrowFlux Universal Dimmer; GrowFlux Access Point, GrowFlux Ltd, Philadelphia, PA, USA) was used. Light intensity target values at canopy level were verified using a PAR-light meter (LI-250A Light Meter and Quantum Photometric Sensor, LI-COR Inc., Lincoln, NE, USA).

Lighting implemented in the small-scale testbed and the plant phenotyping experiments consisted of four lighting regimes, termed ‘Constant-155’, ‘High-Low’, ‘Sunlike’ and ‘Low-High’ (Fig. 1A). All lighting regimes were set to start at 06:00 and consisted of a L18:D6 photoperiod with a cumulative daily light integral (DLI) of 10 mol m^-2^ PPFD. In ‘Constant-155’, an 18-hour photoperiod with light intensity of 155 μmol m^-2^ s^-1^ PPFD was used to emulate a commonly used lighting strategy in leafy vegetable production. In the other three lighting regimes, the 18-hour photoperiod was divided into three 6-hour periods; 0-6 h, 6-12 h and 12-18 h ZT, in which ZT indicates the time elapsed since the onset of photoperiod (from the German-word *zeitgeber* for “time-giver”). During these periods the light intensity was set either to low light (95 μmol m^-2^ s^-1^ PPFD) or high light (275 μmol m^-2^ s^-1^ PPFD) at specific times of the photoperiod. In High-Low lighting regime, high light was applied at 0-6 h ZT, in Sunlike at 6-12 h ZT, and in Low-High at 12-18 h ZT of the photoperiod. During the last 6-hour period, at 18-24 h ZT, the light intensity was set to 0 μmol m^-2^ s^-1^ PPFD (darkness) in all regimes.

**Figure 1.**
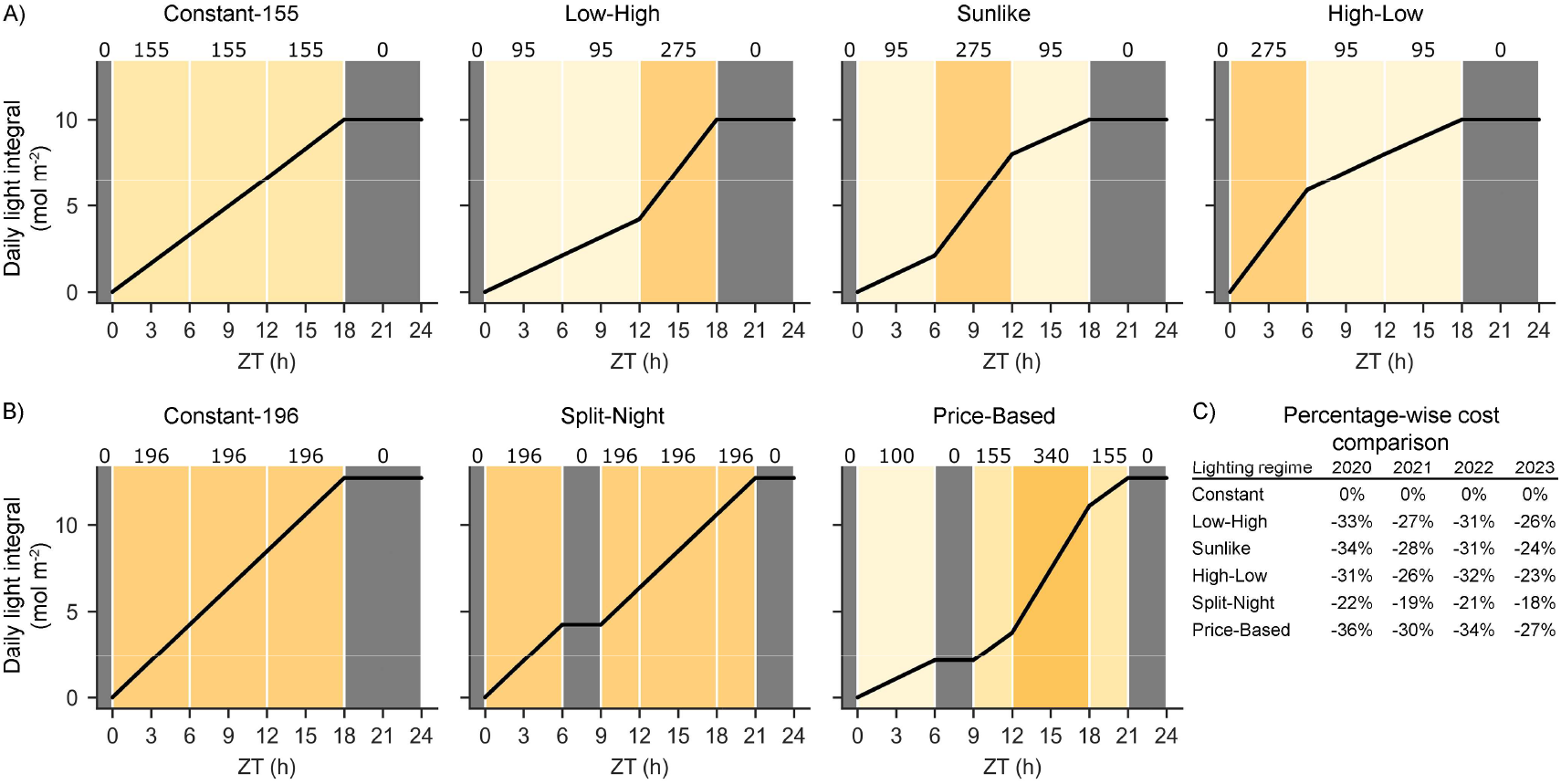
Lighting regimes used in experiments on lettuce cv. ‘Katusa’. A) In a small-scale vertical farm testbed and plant phenotyping platform, the lighting regimes were termed Constant-155, Low-High, Sunlike and High-Low, with an equal daily light integral (DLI) of 10.0 mol m^-2^ PPFD. B) In a larger-scale vertical farming experiment, the lighting regimes were termed Constant-196, Split-Night and Price-Based, with different light intensity levels but an equal DLI of 12.7 mol m^-2^ PPFD. Black lines indicate cumulative daily light integral values, numbers on top of the panels and different shades of yellow in the background indicate target light intensity values (PPFD). ZT (h) indicates time in hours elapsed since the beginning of the photoperiod. C) Percentage-wise comparison of potential electricity cost-savings of dynamic lighting regimes between years 2020-2023 in Finland, with constant lighting regimes used as a baseline (Supplemental table 2).

Lighting in the larger-scale vertical farming experiment consisted of three lighting regimes, termed ‘Constant-196’, ‘Split-Night’ and ‘Price-Based’ (Fig. 1B). All lighting regimes were set to start at 11:00 and consisted of photoperiods with a cumulative DLI of 12.7 mol m^-2^ PPFD. Constant-196 consisted of L18:D6 at 196 μmol m^-2^ s^-1^, Split-Night was L6:D3:L12:D3 with lighting at 196 μmol m^-2^ s^-1^, and Price-Based was L6:D3:L12:D3 with a more complex profile of 6 h of light at 100 μmol m^-2^ s^-1^, 3 h of darkness, 3 h of light at 150 µmol m^-2^ s^-1^, 6 h of light at 340 µmol m^-2^ s^-1^, 3 h of light at 150 µmol m^-2^ s^-1^, and 3 h of darkness. The light intensities in Split-Night and Price-Based followed a pattern that reflected changing electricity market prices during a 24-hour cycle.

The three experimental setups differed with respect to the spectral composition of illumination. In the small-scale vertical farm testbed, we used Valoya AP673L spectrum (BX120, Valoya Greenlux Lighting Solutions Oy, Helsinki, Finland) (Supplemental Fig. 1). The plant phenotyping experiment was conducted under a PSI spectrum (CS 250/300_4_2.4, PSI, Drásov, Czech R.). In the larger-scale experiment, we used PhysioSpec Greenhouse spectrum (VYPR, Fluence Bioengineering Ltd, Austin, Texas, USA) for initial growth and Valoya Solray spectrum (BX120, Valoya Greenlux Lighting Solutions Oy, Helsinki, Finland) (Supplemental Fig. 1). The spectral photon irradiance was measured with an array spectroradiometer (Maya2000 Pro Ocean Optics, Dunedin, Fl, USA; D7-H-SMA cosine diffuser, Bentham Instruments Ltd, Reading, UK) (Supplemental Fig. 1). Measurements were recorded within the wavelength range 315-800 nm and processed in R (R Core Team, 2017), using the photobiology packages developed for spectral analysis (Aphalo, 2015).

### Calculation of potential electricity cost savings

The potential electricity cost savings of dynamic lighting regimes were calculated using the following method: 1. The hourly dynamics of average electricity price fluctuations within a 24-hour period (in €/kWh) were calculated for different timescales (monthly, yearly) using historical electricity spot price data between 2020-2023 in Finland (https://www.nordpoolgroup.com/en/elspot-price-curves/); 2. The average price fluctuation patterns were used to identify, rank, and target the hours of the day which yield the highest reductions in electricity costs; 3. The hourly target light intensity values of the lighting regimes between experiments were scaled to an equal DLI, and then used in conjunction with the corresponding electricity consumption data of the LED-lights to align with the lowest electricity prices; 4. The total electricity costs of different lighting regimes for a 24-hour period were calculated for each hour throughout the photoperiod; 5. Finally, the total electricity costs (€/m^2^) of the different lighting regimes were calculated to allow percentage-wise comparison of potential electricity cost-savings. (Supplemental Table 2).

### Image-based analysis of growth, colour and chlorophyll fluorescence

In the plant phenotyping experiment in NaPPI, visible light RGB imaging was used to record individual plant growth, morphology, and colour under the different lighting regimes. Top view images were captured at 8, 12, 16, 20, 24, and 28 days after sowing (DAS) to record changes in canopy cover area over time. Plantscreen Data Analyzer v3.2.4.5 (PSI, Drásov, Czech R.) was employed to post-process the RGB images. The plant surface area was segmented using pixel colour thresholding for RGB images. Morphological and physiological parameters were extracted automatically by PlantScreen Data Analyzer. Plant canopy area expansion rates (mm^2^ · day) from 12 DAS to 28 DAS were determined using the following equation:

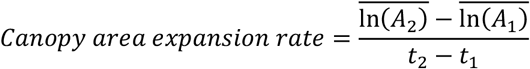

where is 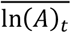 is the averaged natural logarithm-transformed canopy cover area at time *t*; *A*_2_ and *A*_1_ are plant canopy cover areas at *t*_2_ and *t*_1_, respectively; and *t*_2_ and *t*_1_ are harvest days 28 and 12 DAS.

Canopy Greenness was computed from RGB images using Python version 3.10.7 and OpenCV (CV2, https://opencv.org/). Pixel values for Red (R), Green (G), and Blue (B) channels were averaged to obtain the average Red (avgR), Green (avgG), and Blue (avgB) values. The Greenness was then determined using the formula by Signorelli et al. (2023):

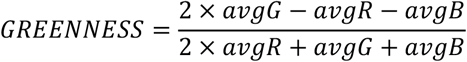

Chlorophyll fluorescence (ChlF) and canopy leaf greenness were analysed between 27-28 DAS, 30 min before the light intensity changes took place at 0 h, 6 h, 12 h, and 18 h ZT. ChlF measurements were performed using a FluorCam FC-800MF pulse amplitude modulated (PAM) chlorophyll fluorometer (PSI, Drásov, Czech R.). ChlF parameters were determined using a quenching protocol developed with FluorCam 7.0 software (PSI, Drásov, Czech R.). Saturating light pulses were applied during the following steps: 1) 20 minutes of dark adaptation, 2) 70 seconds of actinic light, and 3) 100 seconds of dark relaxation. Shutter speed and sensitivity were set to 33 µs and 5%.

In the larger-scale vertical farming experiment, ChlF imaging was performed on the last day of the experiment at a two-hour interval between 0-24 h ZT using the PlantScreen SC Mobile System (PSI, Drásov, Czech R.). The ChlF protocol was executed after a 30-min dark acclimation and included a 9-min high light period (800 μmol m^-2^ s^-1^) followed by a 27-min dark relaxation and then a light ramp consisting of 3-min steps of increasing light intensity (100, 200, 300, 600, 800, 1000 and 1200 μmol m^-2^ s^-1^). Saturating light pulses were triggered every 3 minutes throughout the protocol to determine quantum yields of PSII and NPQ. NPQ was calculated as (Fm-Fm’)/Fm’ where Fm and Fm’ are maximum fluorescence yields in dark- and light-acclimated conditions, accordingly (Horton and Ruban, 1992). Values for NPQ were calculated at nine minutes into the dark relaxation period and termed qErel-D9-min. The ChlF data was post-processed using Fluorcam10 software (PSI, Drásov, Czech R.). Plant segmentation using pixel colour thresholding followed the same principle as for the plant phenotyping experiment, while data extraction was conducted with a custom processing pipeline.

### Analysis of photosynthetic activity and growth

Photosynthetic gas exchange rates (A = µmol CO_2_ m^-2^ s^-1^) in the plant phenotyping experiment were measured at 28 DAS at timepoints 3 h, 9 h and 15 h ZT using LI-6400/XT Portable Photosynthesis System (LI-COR Inc., Lincoln, NE, USA). For each timepoint, 5 replicate plants from each lighting regime were measured. Values for each plant consisted of three technical replicate averages.

In all experimental setups, above-ground plant fresh weights (FW) were measured immediately after harvesting using a digital scale. In the plant phenotyping and larger-scale vertical farming experiment, the samples were oven-dried at +60°C for 7-14 days, after which their dry weights (DW) were measured.

### Analysis of metabolites

For the analysis of soluble carbohydrates and starch in the small-scale vertical farming experiment, samples were harvested at the end of the experiment at a six-hour timepoint interval before the light intensity changes took place at 0 h, 6 h, 12 h, and 18 h ZT. From each lighting regime, 3 individual plants per timepoint were collected. In the larger-scale vertical farming experiment, samples were harvested at the end of the experiment at a two-hour timepoint interval between 0-24 h ZT. From each lighting regime, three replicate sets of 24 leaf disks (diameter 8 mm) from a pool of 4 plants (6 disks per plant) per timepoint were collected. Immediately after harvesting, the samples were placed in vials, flash-frozen in liquid nitrogen and stored in −80°C. Frozen samples were ground to fine powder with a ball-mill (Retsch MM 400 Mixer Mill, Retsch GmbH, Haan, Germany). For the analysis of glucose, fructose, sucrose, starch, chlorophyll, total protein, and nitrate concentrations, metabolites were extracted twice with 80% ethanol and once with 50% ethanol from 20 mg of frozen tissue powder and analysed as described in Cross et al. (2006).

For large-scale metabolite profiling, metabolites were extracted from approximately 50 mg (FW) of plant material per sample as described in Sipari et al (2022) (for details, see Supplemental data 1). Feature extraction from raw LC-MS data was conducted using MZmine (Version 4.0.3) in batch mode and the resulting features were further processed with Python (Version 3.11.3). The features were filtered based on retention time (0.4–4 minutes) and a minimum feature intensity of 10 across all samples. Additionally, features were required to have a relative standard deviation (RSD) of less than 150% across QC samples and to be present in at least two-thirds of the samples within a sample class. Unsupervised dimension reduction was done using principal component analysis (PCA). For PCA, QC samples were excluded from the dataset and the missing values were imputed using the lowest 10% of feature-wise intensity. The dataset was then log-transformed, and Z-score normalized. The normalized data was projected onto the first two principal components (PC1 and PC2).

### Statistical analyses

All three experimental setups were based on randomized complete block design (RCBD), with different lighting regimes regarded as experimental units. Time-independent experiment repetitions, growth chamber shelf units and compartments used in the experiments were regarded as block grouping factors. Individual plant replicates were regarded as subsamples or observational units.

The small-scale vertical farming experiment was repeated twice. Plants in each lighting regime were grown on separate shelves, and each time-independent repetition with 12 and 24 subsamples in block 1 and 2, respectively, was regarded as a block. The plant phenotyping experiment in NaPPI was conducted once. Plants in each lighting regime were grown on two separate shelves, and each shelf unit with 10 subsamples was regarded as a block. The larger-scale vertical farming experiment was repeated three times. Plants in each lighting regime were grown in three separate compartments on a rotating basis between repetitions, and each time-independent repetition with 35-40 subsamples was regarded as a block.

FW data from all three experimental setups, DW data from the plant phenotyping and larger-scale vertical farming experiments, canopy cover area, roundness and compactness from the plant phenotyping experiment, and soluble carbohydrates and chlorophyll fluorescence parameters from the larger-scale vertical farming experiment were analysed by fitting a linear mixed effects model, with random effects for the block/shelf/experiment grouping factors. Analysis was performed with R software (R Core Team, 2024), using package NLME (Pinheiro & Bates, 2000, 2023). In cases when variances were not homogeneous, a power of variance function covariate was included in the model. If the test of overall significance of the treatments yielded *P* < 0.05, comparisons between individual pairs of treatments were done with package GMODELS (Warner et al. 2024) to fit these contrasts. *P*-values from multiple contrasts were adjusted using Holm’s procedure. *P* = 0.05 was used as the limit for significance of treatment effect, while for pairwise comparisons *P* = 0.10 was used. Concentrations of soluble carbohydrates, chlorophylls, total protein, and nitrate from the small and larger-scale vertical farming experiments and photosynthetic gas exchange rates from the plant phenotyping experiment were analysed with analysis of variance (ANOVA) and Tukey HSD post-hoc test by comparing values at each timepoint between lighting regimes.

## Results

### Dynamic lighting regimes support lettuce cv. ‘Katusa’ growth in different indoor cultivation setups

Lettuce cv. *‘*Katusa*’* was first grown under four alternative lighting regimes sharing the same DLI: Constant-155, or dynamic Low-High, Sunlike and High-Low (Fig. 1A) in two different growth environments, a small-scale vertical farm testbed and plant phenotyping platform. Comparison of electricity costs between constant and dynamic lighting regimes indicated up-to −36% potential cost savings in dynamic lighting as compared to constant lighting, when periods of higher light intensity and electricity consumption were aligned with the lowest electricity spot price hours (Fig. 1C, Supplemental Table 2).

In the plant phenotyping platform, no significant differences were observed in plant fresh weight at the end of the experiment 23 DAS (Table 1). However, based on above-ground dry matter content (%), growth analysis showed small but statistically significant differences between constant vs. dynamic lighting (Table 1). Plants grown under Constant-155 accumulated approximately 7.14 % dry matter, whereas plants grown under Low-High, High-Low and Sunlike regimes accumulated 6.56 %, 6.73 % and 6.44 % dry matter, respectively (Table 1). Analysis of fresh weight in the small-scale vertical farm revealed small but statistically significant differences 29 DAS, with highest value of 6.13 g FW in Constant-155, followed by High-low with 5.78 g FW, Low-high with 5.31 g FW, and Sunlike with 5.24 g FW (Table 1).

**Table 1.**
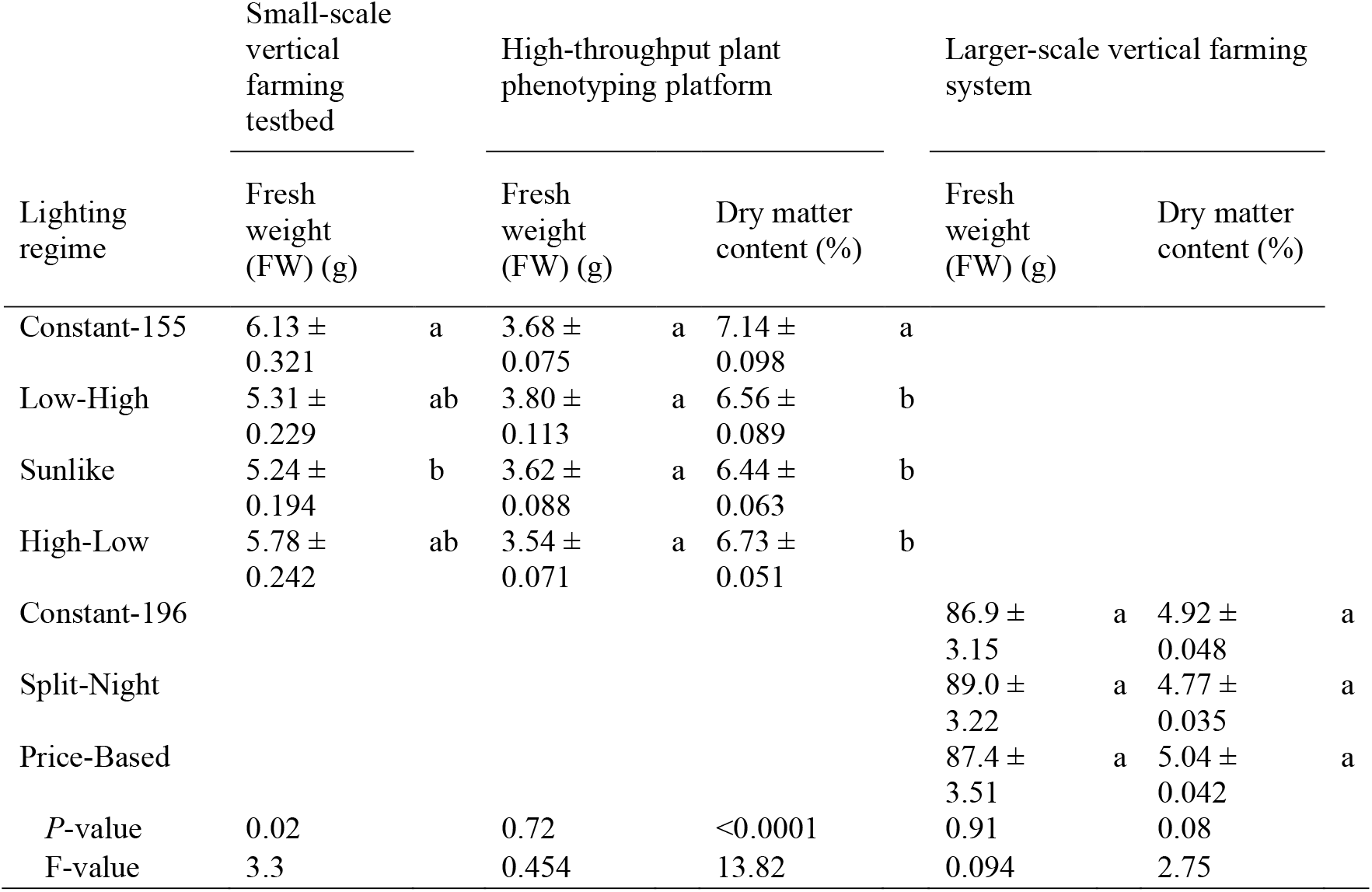
Analysis of lettuce cv. ‘Katusa’ fresh weight (g) and dry matter content (%) in the small-scale vertical farming testbed (n=36, df=3 and 139) and high-throughput plant phenotyping platform (n=20, df=3 and 61) under Constant-155, Low-High, Sunlike and High-Low lighting regimes, and in the larger-scale vertical farming system (n=115-116, df=2 and 31) under Constant-196, Split-Night and Price-based lighting regimes. Data represent mean ± SE. Letters indicate statistical significance between treatments (*P* < 0.05).

In the plant phenotyping platform, seedling growth was further assessed by RGB imaging of canopy cover area at a 4-day interval between 12 DAS and 28 DAS (Table 2). Canopy cover area expansion rates (mm^2^ day^-1^) were observed to be higher in Low-High, Sunlike and High-Low than in Constant-155 (Table 2). However, plant morphological characteristics measured as canopy roundness and compactness were not affected by the lighting regimes (Table 2).

**Table 2.** Analysis of lettuce cv. ‘Katusa’ morphological parameters in the plant phenotyping platform (n=20) under Constant-155, Low-High, Sunlike and High-Low lighting regimes. Canopy cover area treatment df=3 and 61 and roundness, compactness, and canopy cover expansion rate treatment df=3 and 75. Data represent mean ± SE. Letters indicate statistical significance between treatments (*P* < 0.05).

| Lighting regime | Roundness |  | Compactness |  | Canopy cover area (mm<sup>2</sup>) |  | Canopy cover area expansion rate (mm<sup>2</sup> day<sup>-1</sup>) |  |
| --- | --- | --- | --- | --- | --- | --- | --- | --- |
| Constant-155 | 0.390 ± 0.012 | a | 0.840 ± 0.005 | a | 6273 ± 118 | a | 0.204 ± 0.009 | a |
| Low-High | 0.380 ± 0.012 | a | 0.837 ± 0.006 | a | 6863 ± 127 | a | 0.215 ± 0.011 | b |
| Sunlike | 0.410 ± 0.008 | a | 0.851 ± 0.004 | a | 6653 ± 152 | a | 0.215 ± 0.009 | b |
| High-Low | 0.380 ± 0.010 | a | 0.833 ± 0.005 | a | 6402 ± 115 | a | 0.210 ± 0.010 | ab |
| *P*-value | 0.15 |  | 0.082 |  | 0.24 |  | 0.0014 |  |
| F-value | 1.81 |  | 2.32 |  | 1.43 |  | 5.75 |  |

Next, we designed more complex lighting regimes to follow lettuce growth under conditions which could be scaled up to commercial production. These lighting regimes were tested in a larger-scale vertical farming system and included Constant-196, Split-Night and Price-Based regimes (Fig. 1). The latter two were characterized by a bipartite split-night photoperiod that reflected the changes in electricity price over a 24-hour cycle (Fig. 1B). Both Split-Night and Price-Based lighting regimes included two 3-hour periods of darkness, which were designed to coincide with the distinct peak hours in demand and electricity price that typically occur in the early morning and afternoon/early evening. Calculation of energy costs for the Split-Night regime indicated up to −22% cost saving, while the Price-Based regime conferred up-to −36% saving in electricity costs (Fig. 1C, Supplemental Table 2). The impact of these lighting regimes on lettuce growth was examined in experiments in which the plants were grown up to 35 days to reach a larger head size, which represents saleable sizes in commercial lettuce cultivation. Neither Split-Night nor Price-Based affected lettuce growth when compared to growth under constant light, as indicated by comparable fresh weights and dry matter contents observed at harvest (Table 1). Flowering was not induced in any of the conditions studied. These findings suggested bipartite split-night regimes with dynamic light intensities as a promising tool to optimize resource use efficiency in lettuce production.

### Lettuce cv. ‘Katusa’ displays photosynthetic adjustments that follow the intensity and timing of dynamic lighting

The functional status of photosynthetic light reactions was followed in the NaPPI phenotyping platform that enabled manual photosynthetic gas exchange measurements as well as automated RGB and chlorophyll *a* fluorescence imaging under Constant-155, Low-High, Sunlike and High-Low lighting regimes. Photosynthetic gas exchange was recorded across the four lighting regimes 3, 9, and 15 hours into the photoperiod, i.e. in the middle of each 6-hour lighting period (Fig. 2A, Supplemental table 4). Highest photosynthetic activities were consistently observed during each high light illumination period in Low-High, Sunlike and High-Low. Lettuce cv. ‘Katusa’ grown under Constant-155, Sunlike and High-Low lighting regimes displayed a typical response with end of the day repression of photosynthetic carbon assimilation at the 15-hour timepoint (Fig. 2A, Supplemental table 4). Plants grown under Low-High lighting regime, however, displayed increased photosynthetic activity at the latest timepoint (15 h) (Fig. 2A, Supplemental table 4).

**Figure 2.**
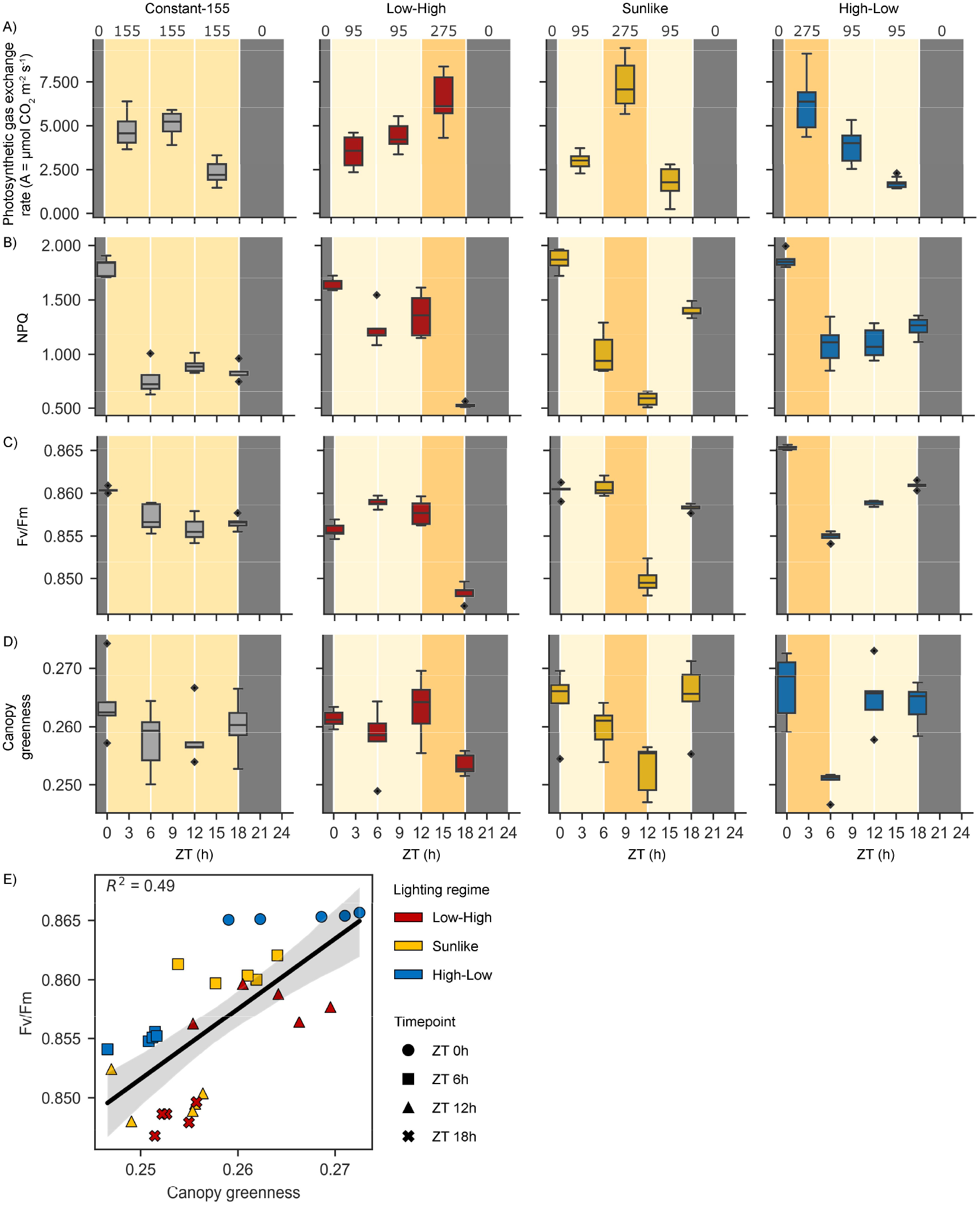
Dynamic regulation of photosynthetic traits in lettuce cv. ‘Katusa’ grown under Constant-155, Low-High, Sunlike and High-Low regimes in the plant phenotyping platform. Numbers on top of the panels and different shades of yellow in the background indicate target light intensity values (PPFD). ZT (h) indicates time in hours elapsed since the beginning of the photoperiod. A) Photosynthetic gas exchange rates (A = µmol CO_2_ m^-2^ s^-1^) were measured at three timepoints, in the middle of each 6-hour lighting period at 3 h, 9 h and 15 h ZT. B) NPQ, C) Fv/Fm and D) canopy greenness index was measured at four timepoints, 30 minutes before light intensity changes took place at 0 h, 6 h, 12 h and 18 h ZT. E) Correlation of Fv/Fm and canopy greenness between timepoints preceding and following a high light illumination period in lighting regimes Low-High, Sunlike and High-Low. Symbol shapes and colours indicate measurement timepoints and lighting regimes, respectively. The photosynthetic parameters were analysed from five plants (n=5) per lighting regime and timepoint. Statistical data analysis by analysis of variance (ANOVA) and Tukey HSD post-hoc test is presented in Supplemental tables 3 and 4.

The RGB and chlorophyll *a* fluorescence images were obtained 30 min before the light intensity shifts took place at the end of each of the 6-hour lighting periods at 0, 6, 12, and 18 hours into the photoperiod (Fig. 2B-D, Supplemental table 3). Under all dynamic lighting regimes, a small but consistent decrease was observed in NPQ, Fv/Fm and canopy greenness index at the end of a 6-hour high light period, regardless of the time of day (Figs 2B-D, Supplemental table 3). Moreover, the analysis of correlation between Fv/Fm and canopy greenness index upon transfer from low to high light intensity revealed that different lighting regimes formed distinct clusters (Fig. 2E). This suggested that the dynamics of leaf greenness and photosynthetic efficiency were interconnected and sensitive to the lighting regime.

### Quantum efficiency of photosynthesis and NPQ respond dynamically to alternative lighting regimes

For more detailed analysis of the effects of dynamic lighting regimes on photosynthetic light reactions, chlorophyll *a* fluorescence imaging was performed every 2 hours across the 24-hour period in the larger-scale vertical farming system. This approach revealed similar pattern of fluctuation in Fv/Fm in the Constant-196 lighting conditions (Fig. 3A, Supplemental table 5). Another parameter following a similar pattern was the partially relaxed NPQ that was calculated at nine min into the dark relaxation period and termed qErel-D9-min (Fig. 3B, Supplemental table 5). Fluctuations in Fv/Fm and qErel-D9-min were also observed under the bipartite Split-Night lighting regime, but the pattern of fluctuation differed from that of the Constant-196 plants (Fig. 3, Supplemental table 5). Under Split-Night conditions, Fv/Fm and qErel-D9-min increased during the first 3-hour dark period but declined upon shift to light (Fig. 3). In contrast, in the Price-Based lighting regime, fluctuations in Fv/Fm and qErel-D9-min were barely detectable (Fig. 3). These results corroborated the finding that the photosynthetic light reactions can delicately adjust their function according to the lighting regime, without detrimental effects on lettuce growth and development (Tables 1-2).

**Figure 3.**
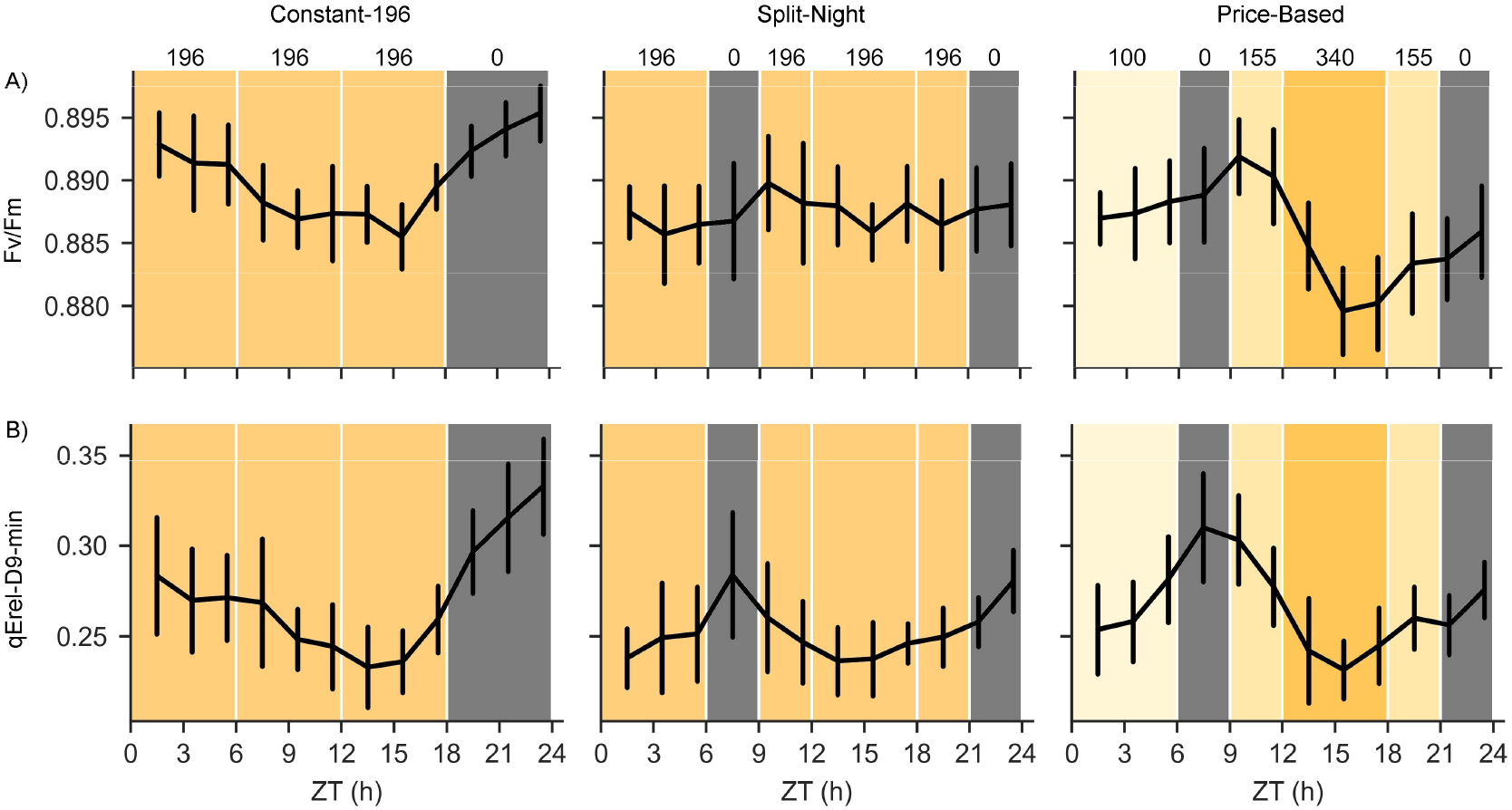
Dynamic regulation of photosynthetic traits in lettuce cv. ‘Katusa’ measured at 2-hour intervals throughout the photoperiod in plants grown under Constant-196, Split-Night and Price-Based lighting regimes in the larger-scale vertical farming system. Numbers on top of the panels and different shades of yellow in the background indicate target light intensity values (PPFD). ZT (h) indicates time in hours elapsed since the beginning of the photoperiod. A) Fv/Fm and B) qErel_D9-min were imaged using a ChlF protocol including a 9-min high light period at 800 μmol m^-2^ s^-1^ PPFD, followed by a 27-min dark relaxation and then a light ramp consisting of 3-min steps of increasing light intensity (100, 200, 300, 600, 800, 1000 and 1200 μmol m^-2^ s^-1^ PPFD). Saturating light pulses were triggered every three minutes throughout the protocol to determine Fv/Fm and NPQ. Values for NPQ were calculated at nine minutes into the dark relaxation period and termed qErel-D9-min. The photosynthetic parameters were analysed from three time-independent experiments including twelve plants (n=12) per lighting regime and timepoint. Statistical data analysis by a linear mixed effects model with random effects for grouping factors is presented in Supplemental table 5.

### Accumulation and depletion patterns of sucrose and starch follow the cumulative daily light integral but the overall metabolic composition of lettuce cv. ‘Katusa’ leaves remains unaltered

To examine the photosynthetic production potential underlying growth under dynamic lighting, the accumulation and depletion patterns of sucrose and starch were analysed across the 24-hour light/dark cycle. In plants grown under Constant-155 in the small-scale vertical farm testbed, the levels of sucrose and starch gradually increased until the end of the 18-hour photoperiod and decreased during the subsequent dark period (Fig. 4, Supplemental table 6). Under Low-High regime, sucrose and starch contents increased slowly during the first 12 hours of the photoperiod and a clear increase was detected during the subsequent high light illumination period (Fig. 4, Supplemental table 6). In Sunlike conditions, the levels of sucrose and starch increased during the high light period applied at midday (Fig. 4, Supplemental table 6). In High-Low, sucrose and starch contents increased rapidly during the early high light illumination period but the accumulation slowed down over the rest of the photoperiod in low light intensity (Fig. 4, Supplemental table 6). In all lighting regimes, the patterns of sucrose and starch contents were associated with the cumulative daily light integral, reaching similar concentration levels at the end of photoperiod (Fig. 1A, Fig. 4, Supplemental table 6). During the night, the levels of both sucrose and starch decreased. Parallel measurements of fructose, glucose, chlorophyll, total protein, and nitrate contents did not reveal light quantity-dependent patterns during the photoperiod (Supplemental table 6). Likewise, in the larger-scale vertical farming experiment, the sucrose and starch contents followed changes occurring in the light intensity, measured under Constant-196, Split-Night and Price-Based strategies every 2 hours during a 24-hour photoperiod (Fig. 5, Supplemental table 7).

**Figure 4.**
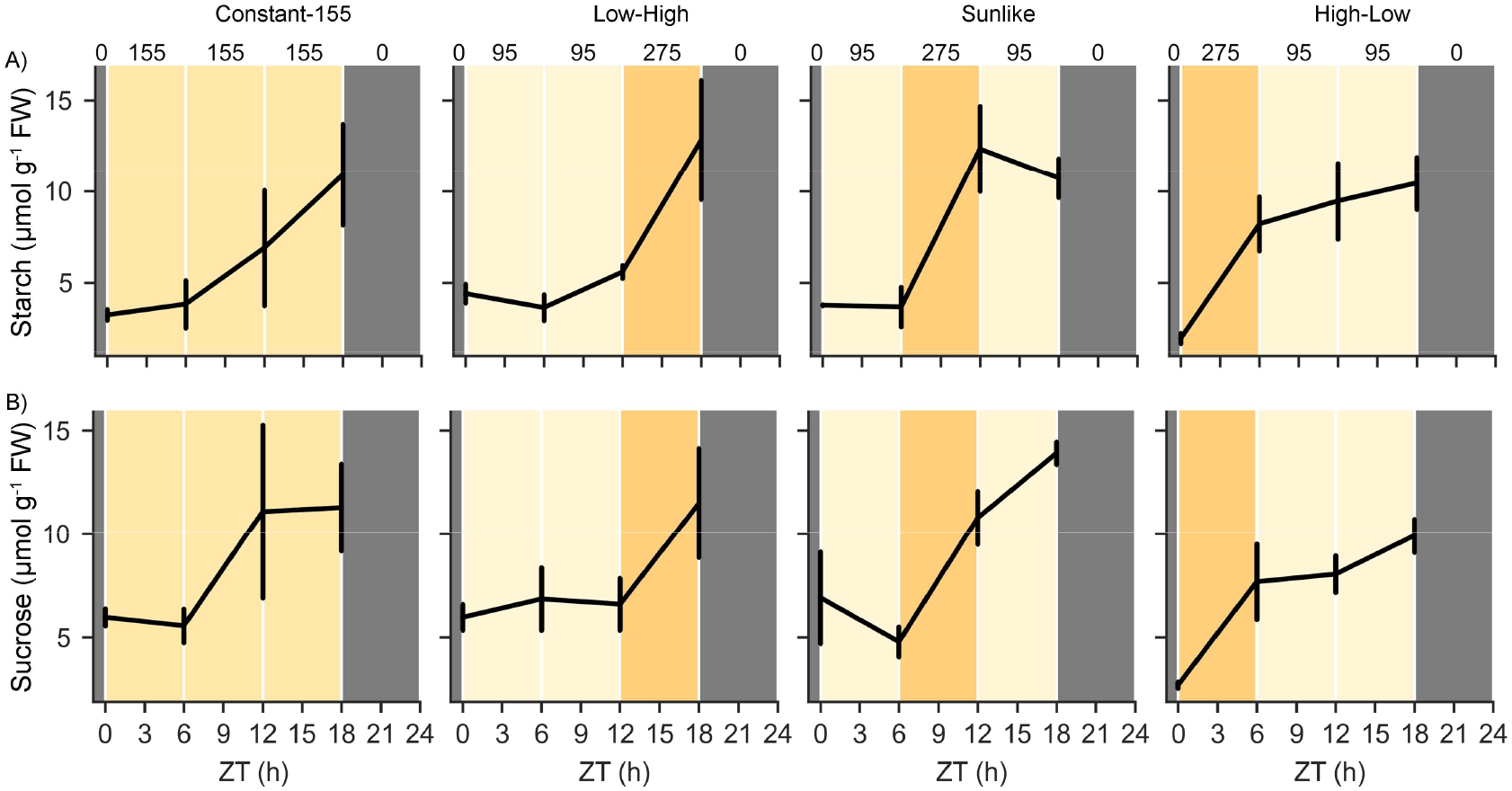
Contents of A) starch and B) sucrose (µmol g^-1^ FW) in lettuce cv. ‘Katusa’ measured at timepoints ZT 0 h, 6 h, 12 h and 18 h in plants grown under Constant-155, Low-High, Sunlike and High-Low regimes in the small-scale vertical farm testbed. Numbers on top of the panels and different shades of yellow in the background indicate target light intensity values (PPFD). ZT (h) indicates time in hours elapsed since the beginning of the photoperiod. Starch and sucrose contents were analysed from three plants (n=3) per lighting regime and timepoint. Statistical data analysis of analysis of variance (ANOVA) and Tukey HSD post-hoc test is presented in Supplemental table 6.

**Figure 5.**
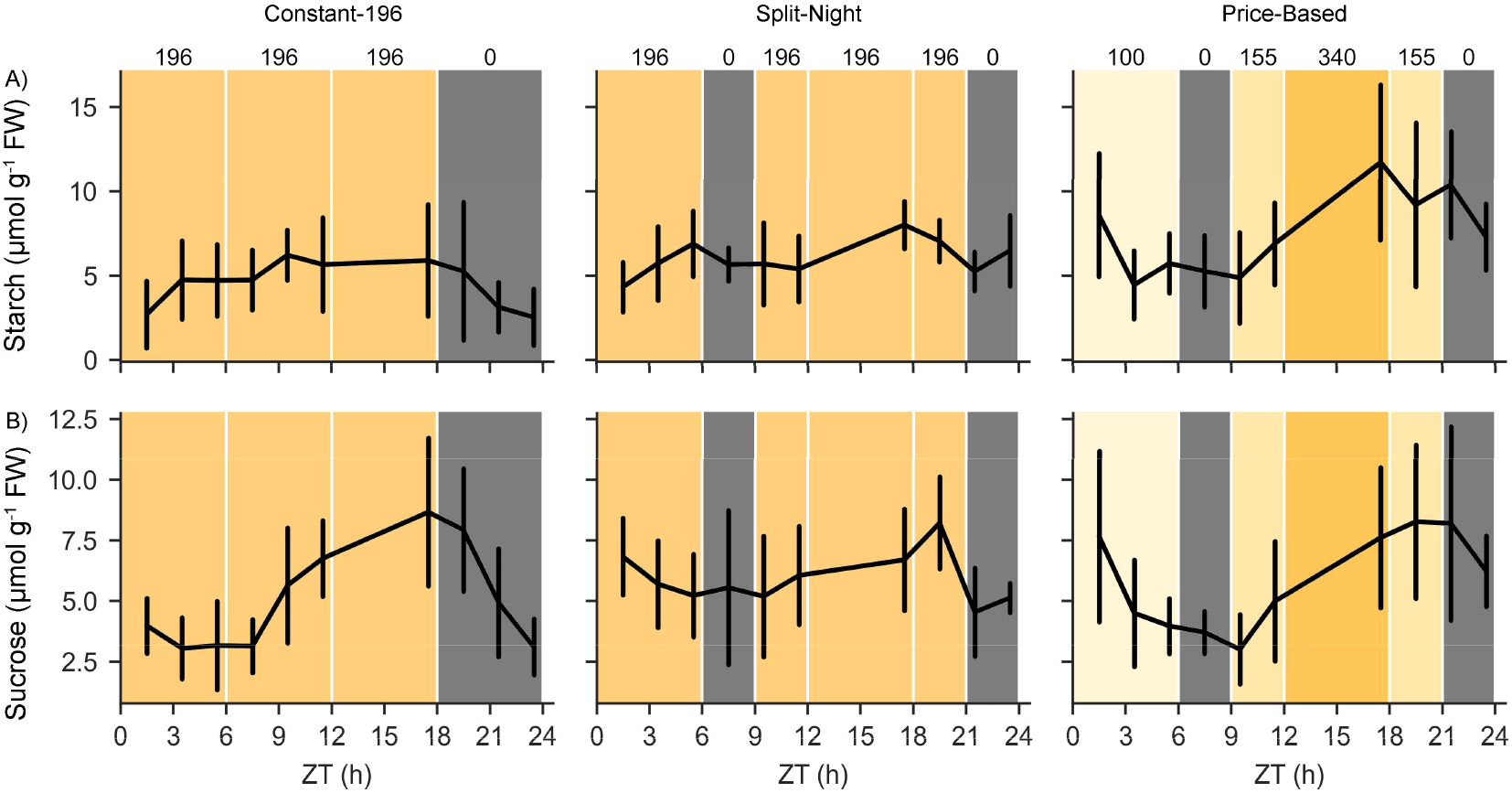
Contents of A) starch and B) sucrose (µmol g^-1^ FW) in lettuce cv. ‘Katusa’ measured at 2-hour intervals throughout the photoperiod in plants grown under Constant-196, Split-Night and Price-Based regimes in the larger-scale vertical farming system. Numbers on top of the panels and different shades of yellow in the background indicate target light intensity values (PPFD). ZT (h) indicates time in hours elapsed since the beginning of the photoperiod. Starch and sucrose contents were analysed from three time-independent experiments including nine biological replicates (n=9) per lighting regime and timepoint. Statistical data analysis of a linear mixed effects model with random effects for grouping factors is presented in Supplemental table 7.

Finally, untargeted metabolite profiling across the 24-hour light/dark cycle was conducted to examine whether growth under dynamic lighting regimes induced wider metabolic shifts in plants grown in the small-scale vertical farming testbed. Altogether 199 compounds were detected using negative mode and 271 compounds using positive mode in MS analysis (Supplemental data 1). The identified compounds included aromatic amino acids, panthotenic acid (vitamin B5), various phenolic compounds including caffeic acid, quercetin and kaempherol derivatives, and sesquiterpenoid lactones (Supplemental data 1). Multivariate analysis of metabolite profiles did not reveal clustering among sample sets (Fig. 6). Rather, the slight variations between individual samples suggesting that growth under moderately alternating dynamic lighting did not deteriorate the metabolite content of lettuce cv. ‘Katusa’ leaves (Fig. 6).

**Figure 6.**
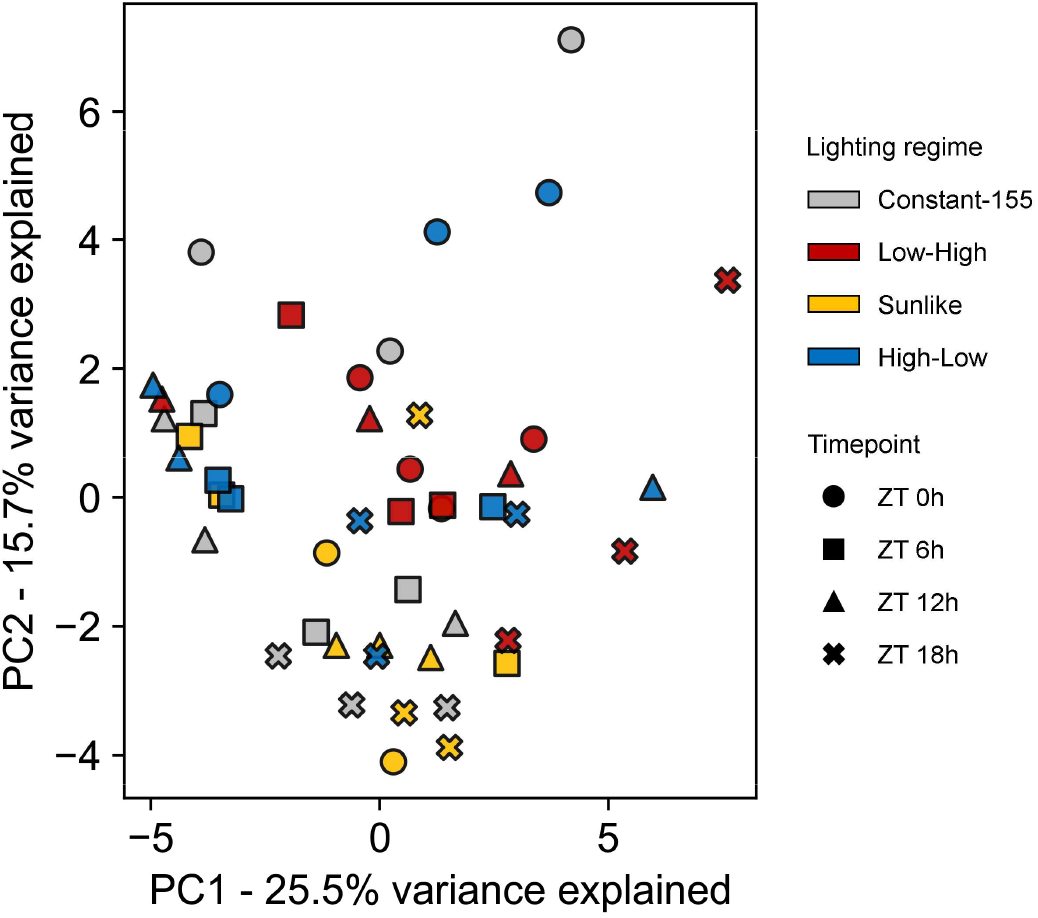
Principal component analysis (PCA) of metabolite profiles in lettuce cv. ‘Katusa’ grown under Constant-155, Low-High, Sunlike and High-Low regimes in the small-scale vertical farm testbed. The samples were collected 15 minutes before light intensity changes took place at timepoints ZT 0 h, 6 h, 12 h and 18 h including a pool of three plants (n=3) per lighting regime and timepoint. Symbol colours and shapes indicate lighting regimes and measurement timepoints, respectively.

## Discussion

Dynamic LED-based lighting technologies could forward controlled environment agriculture by enabling cost-effective growth without compromising the yield or nutritional quality of the produce. The linear relationship between light intensity and electricity consumption of LED luminaires can directly be utilized in reducing electricity-related production costs. Here, we tested various dynamic lighting regimes where the intensity of artificial lighting was altered at specific times of the photoperiod, mirroring average changes in electricity price. Estimation of electricity costs using average spot-price information from NordPool for 2020-2023 indicated up-to −36% cost savings under dynamic lighting (Fig. 1, Supplemental Table 2). High-throughput plant phenotyping techniques and growing conditions simulating vertical farming production systems were utilized to assess lettuce cultivation. We found that lettuce cv. ‘Katusa’ adjusted its photosynthetic processes dynamically according to the prevailing light intensity throughout the photoperiod (Figs 2-5), enabling growth and productivity under various dynamic lighting regimes (Tables 1 and 2). Lettuce is among the most cultivated leafy vegetables in CEA and comprises a feasible model plant for physiological studies. Future research should integrate data from different crop species and genetically distinct cultivars to uncover the molecular mechanisms underlying tolerance to dynamic lighting among horticultural crops.

### Dynamic lighting conditions maintain lettuce growth in different indoor cultivation setups

We assessed lettuce growth in three different setups, including a plant phenotyping platform, a vertical farm testbed, and a larger scale vertical farming system with different lighting regimes and spectral qualities, but similar DLI within each indoor farming setup (Fig. 1). The applied dynamic lighting regimes could reduce LED lighting-related electricity costs by up to −36%, when calculated by scheduling the high light episodes to periods of lowest average electricity market prices (Supplemental Table 2). This estimation is in line with studies that modelled and experimentally validated potential electricity cost savings of shifting LED light use towards lower demand hours (Avgoustaki and Xydis, 2021, Arabzadeh et al., 2023). Successfully maintaining crop production in such scenarios requires knowledge on plant growth and development as well as technical energy system concepts (Arabzadeh et al., 2023).

In our experiments, the shifting light intensity periods were programmed to occur repeatedly at specific times of the photoperiod, and the magnitude of differences between low and high light intensities was kept at a maximum of 3.4-fold to avoid light-induced stress (Figs 1-3). In these conditions, lettuce cv. ‘Katusa’ grew comparatively well both under constant and dynamic lighting regimes, indicating that DLI can be redistributed across the photoperiod in different ways (Tables 1 and 2). These findings were in line with Poorter et al., (2019), who conducted a larger-scale meta-analysis of 500 experiments encompassing 760 plant species and found that physiological, developmental, and chemical plant traits were all responsive to the DLI.

Understanding how changing light conditions affect growth in different species is key to optimizing growth conditions for commercial production. Earlier studies have suggested that drastic changes in light conditions could impede plant growth (Morales and Kaiser, 2020, Bhuiyan and van Iersel, 2021). Studies on *A. thaliana* models, aimed at identification of photoprotective components, were often designed to cause light stress with periods of low and high light alternating on a time scale of minutes (Suorsa et al, 2012, Kono et al, 2017, Garcia-Molina and Leister 2020). The outcomes of laboratory studies suggested that dynamic acclimation to changing light intensities involves regulatory interactions among photosynthetic reactions, photoprotective mechanisms, leaf development, and diurnal physiological processes (Schneider et al., 2019, Niu et al., 2024, Shikanai, 2024), which should match the changing light intensities to maintain growth. Recently, Kaiser et al. (2024) tested different leafy vegetables and herbs, including basil (*Ocimum basilicum*), pak choi (*Brassica rapa* subsp. *chinensis*), rucola (*Diplotaxis tenuifolia*), and spinach (*Spinacia oleracea*) under hourly alterations in light intensities, and showed that their marketable fresh weights were not affected, when compared to plants grown under constant light conditions of equal DLI.

We took advantage of a phenotyping platform equipped with adjustable LED luminaires and imaging systems to validate lettuce growth under dynamic lighting. Lettuce displayed similar developmental responses with no undesirable morphological alterations under dynamic vs. constant lighting regimes, even though light intensities changed at different times of the day (Tables 1 and 2). Likewise, Bochenek and Fällström (2016) and Bhuiyan and van Iersel (2021) found that temporal differences in distribution of the DLI did not cause growth defects in biomass accumulation of lettuce cv. ‘Galiano’, ‘Little Gem’ or ‘Green Salad Bowl’. In a larger-scale vertical farming system, leaf biomass accumulation under dynamic lighting did not differ from constant light controls even if the photoperiods were interrupted by two periods of darkness, as demonstrated by lettuce growth under Split-Night and Price-Based lighting regimes (Fig. 1B, Table 1). These conditions, designed to simulate energy-efficient dynamic lighting in commercial production, supported lettuce growth (Table 1) but did not induce flowering in a larger-scale vertical farming system, suggesting scalability of the dynamic lighting strategies.

In our experiments, shifts between light and dark as well as transitions between light intensity levels during the photoperiod were instantaneous and abrupt (Fig. 1). In natural lighting conditions, however, diurnal changes in light intensity occur gradually over time in timescales varying from minutes to hours. Twilight length, or the duration of light intensity transitions at dawn and dusk has been found to influence plant growth and flowering time in Arabidopsis via the *LHY/CCA1*-circadian clock pathway (Mehta et al., 2024). Plants grown under photoperiods with twilight lengths between 30 and 60-minutes grew larger than plants in no-twilight photoperiods (Mehta et al., 2024). As precise control of light intensity can be achieved with LED-luminaires, further studies should examine how gradual transitions in light intensity influence plant development and growth in CEA.

### Photosynthetic metabolism responds to dynamically changing lighting, while the overall leaf chemical composition does not show short-term fluctuations in response to light

We utilized chlorophyll fluorescence imaging in combination with analysis of photosynthetic gas exchange and metabolite profiling to address the physiological and biochemical responses underlying lettuce cv. ‘Katusa’ growth under dynamic lighting. Additionally, visible light RGB imaging was used to record leaf tissue colour responses during the dynamic light intensity changes. By comparing photosynthetic responses under light intensities changing in a timescale of hours, we found that photosynthetic light reactions and carbon metabolism were delicately balanced according to the duration and intensity of dynamic lighting in lettuce cv. ‘Katusa’ (Figs 2-4, Supplemental tables 3-7).

In our dynamic lighting conditions, the rate of photosynthetic carbon assimilation increased during high light illumination in Low-High, Sunlike and High-Low regimes (Fig. 2A, Supplemental table 4). This was accompanied by decreased NPQ values, suggesting photochemical quenching of chlorophyll fluorescence upon activation of carbon assimilation, which could drain reducing equivalents from the photosynthetic electron transfer chain and thereby alleviate excitation pressure at PSII (Fig. 2). The flexible adjustments in photosynthetic reactions were accompanied by accumulation of starch and sucrose, which followed the quantity of photons received during the day (Fig. 4). Photobiological studies have suggested that plants undergo different photosynthetic adjustments to fluctuating light conditions, depending on the frequency, duration, and intensity of fluctuating light (von Bismarck et al 2023, Vialet-Chabrand et al 2017, Alter et al 2012, Yin and Johnson 2000, Lazzarin et al, 2024). Our dynamic lighting conditions comprised one daily high light period with up-to 3.4-fold differences in light intensities (Fig. 1). These dynamic lighting conditions did not result in decreased photosynthetic productivity when compared to constant light with equal DLI (Figs 1-5), suggesting that metabolic interactions were sufficient to allow dynamic acclimation according to the prevailing lighting.

Following the functional status of photosynthetic light reactions with higher temporal resolution in the larger-scale vertical farming system revealed delicate fluctuations in photosynthetic parameters, which could be detected by chlorophyll fluorescence imaging under the more complex, alternative day/night regimes, such as the Split-Night and Price-Based lighting regimes (Figs 1 and 3, Supplemental table 5). The rapidity of photosynthetic adjustments is important, since delays in light-induced adjustments of photosynthetic gas exchange, carbon assimilation, or balancing of the photosynthetic light reactions could negatively affect photosynthetic activity and yield (Kromdijk et al., 2016, Taylor and Long, 2017). A well-known example is slow relaxation of NPQ upon shift from high to low light intensity, which prolongs the dissipation of excitation energy, thereby limiting the conversion of light into chemical energy needed for photosynthesis (Kromdijk et al., 2016, Niu et al., 2023). Excessive dissipation of excitation energy via NPQ could equally well limit the availability of light energy for photochemistry, thereby limiting resource allocation for growth and development. Our findings suggest that the photosynthetic regulatory circuits of lettuce can cope with light intensity changes introduced on a time-scale of hours, without detrimental effects on accumulation of leaf biomass (Fig. 3, Table 1). Alternative, artificial day/night regimes could therefore offer tools to modulate photosynthetic limitations to plant production.

Comparison of metabolite profiles between plants grown under constant moderate light vs. high light has revealed high-light-induced metabolic responses in model plants and horticultural crop species (Jänkänpää et al., 2012, Ishihara et al., 2024). Changes in primary metabolism as well as accumulation of phenolic compounds, notably anthocyanins and flavonoids, are important responses that can protect plants against potentially damaging effects of light. It is noteworthy that the metabolic responses observed upon growth of plants under constant moderate or high light may stem from the differential DLI received by the plants (Ishihara et al, 2024). In our experiments, lettuce cv. ‘Katusa’ grown under varying light regimes, but equal DLI, displayed photosynthetic adjustments that followed the cumulative DLI, whereas overall metabolite profiles did not display dynamic light-induced alterations during the photoperiod (Fig. 6). Overall, the results of this study suggested that application of dynamic lighting regimes can support lettuce growth without impairing leaf chemical composition. Understanding the dynamics of plant metabolism and overcoming photosynthetic limitations to plant production under changing light intensities are keys to cost-effective indoor cultivation.

## Supplementary data

**Supplemental figure 1**. Spectral distributions and photon flux density per wavelength area in different indoor cultivation setups.

**Supplemental Table 1**. Growing conditions and indoor cultivation setups used in this study.

**Supplemental Table 2**. Electricity cost comparison in the different lighting regimes used in this study.

**Supplemental Table 3**. Statistical analysis of data presented in Figure 2, depicting changes in chlorophyll fluorescence and canopy greenness in plants grown in a plant phenotyping facility.

**Supplemental Table 4**. Statistical analysis of data presented in Figure 2, depicting changes occurring in photosynthetic gas exchange rate in plants grown in a plant phenotyping facility.

**Supplemental Table 5**. Statistical analysis of data presented in Figure 3, depicting changes in chlorophyll fluorescence in plants grown in a larger-scale vertical farm.

**Supplemental Table 6**. Analysis of photosynthetic end products, chlorophyll and nitrate contents in lettuce cv. ‘Katusa’ grown under Constant-155, Low-High, Sunlike and High-Low lighting regimes in the small-scale vertical farm testbed.

**Supplemental Table 7**. Analysis of photosynthetic end products in lettuce cv. ‘Katusa’ grown under Constant-196, Split-Night and Price-Based regimes in the larger-scale vertical farm.

**Supplemental Data 1**. Large-scale metabolite profiling of lettuce cv. ‘Katusa’ grown under dynamic lighting regimes.

## Acknowledgements

We thank Ema Marmara, Juha Näkkilä, Satu Engström, Minna Kavander, Päivi Tuomola and Arto Nieminen for their excellent technical assistance. This work was financially supported by Maiju and Yrjö Rikala Horticultural Foundation, Jenny and Antti Wihuri Foundation, Novo Nordisk Plant Science, Agriculture and Food Biotechnology - Project Grants 2020 (NNF20OC0065026), Academy of Finland (343527, 346140), University of Helsinki Faculty of Agriculture and Forestry and Faculty of Biological and Environmental Sciences, Doctoral Programme in Plant Sciences (DPPS), Viikki Plant Science Centre (ViPS), Finnish Ministry of Agriculture and Forestry, the Research and Innovation Program Fund “Catch the carbon”, decision number VN/28558/2020.

## Author contributions

Conceptualization, AM, AS, TK, SK, PE, SP; Methodologies, AM, AS, SP, HI, NS ; Software/VF design and set up, AM; TK; MP; Investigation, AM, SP, HI, AS, TK, IV ; Validation, AM, SP, AS, NS, JH; Formal analysis, NS, JH, SP; Data curation, AM, NS, JH, SP, TK; Visualization, SP; Writing - original draft, AM, AS, SP, NS, TK, SK; Writing - review & editing, AM, SK, TK, PE, KH; Supervision, SK, PE; Project administration, SK; Resources, SK, KH, MP; Funding acquisition, SK, PE, AM, MP.

## Data availability statement

Data that supports the findings of this study are available in the supplementary material of this article.

## Abbreviations

CEA: Controlled environment agriculture
Cytbf: cytochrome b6f complex
DAS: days after sowing
DLI: daily light integral
LED: light-emitting diode
NPQ: non-photochemica quenching
PSI: Photosystem I
PSII: Photosystem II

