## Supplementary figure 1 for "Photosynthetic adjustments maintain lettuce growth under dynamically changing lighting in controlled indoor farming setups"

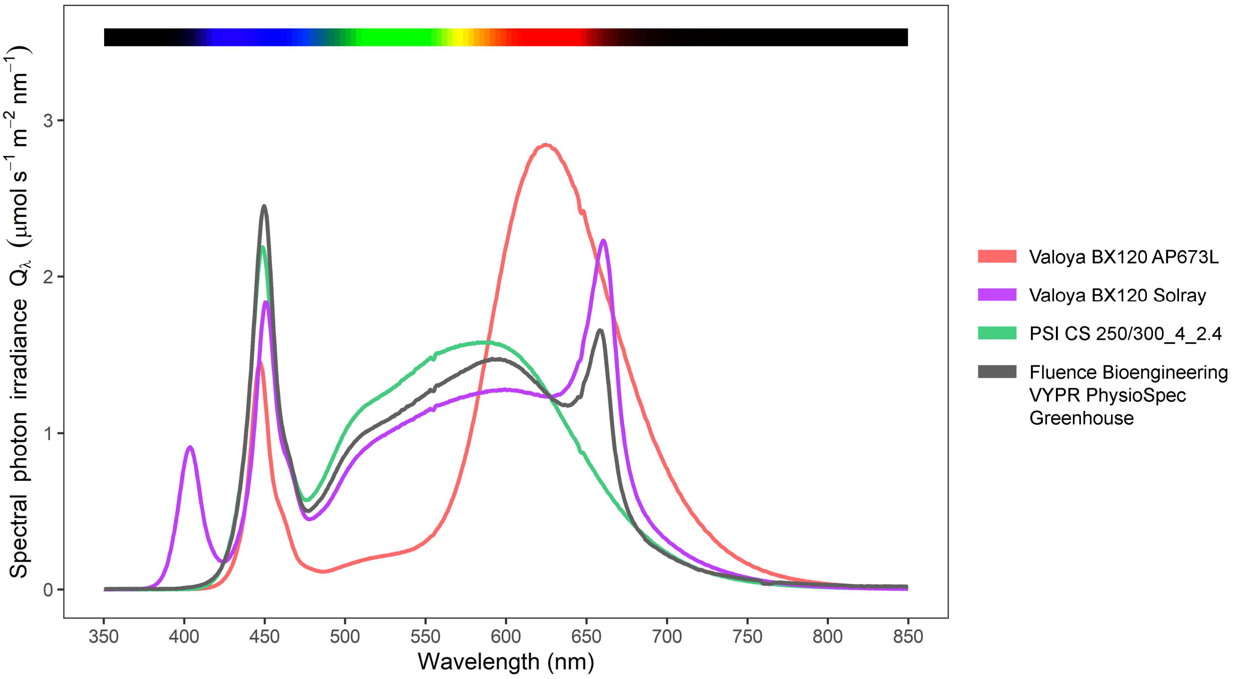


**Supplementary figure 1.** Spectral distributions and photon flux densities per wavelength area in different indoor cultivation setups. Plant phenotyping experiments were conducted under a cool white spectrum (PSI spectrum CS 250/300_4_2.4) (green line). In a small-scale vertical farming testbed, Valoya BX 120 AP673L with a spectrum designed for commercial leafy vegetable cultivation was used (red line). In a large-scale vertical farming experiment, plants were initially grown under Fluence Bioengineering VYPR PhysioSpec Greenhouse LED lighting in a standard greenhouse (gray line) and thereafter transferred to a large-scale vertical farming system equipped with Valoya Solray spectrum BX120 designed for commercial leafy vegetable cultivation (violet line).
